# Venom tradeoff shapes interspecific interactions, physiology and reproduction

**DOI:** 10.1101/2023.07.24.550294

**Authors:** Joachim M. Surm, Sydney Birch, Jason Macrander, Adrian Jaimes-Becerra, Arie Fridrich, Reuven Aharoni, Rotem Rozenblat, Julia Sharabany, Lior Appelbaum, Adam M. Reitzel, Yehu Moran

**Affiliations:** Department of Ecology, Evolution and Behavior, Alexander Silberman Institute of Life Sciences, Faculty of Science, The Hebrew University of Jerusalem, Jerusalem, Israel; Department of Biological Sciences, University of North Carolina at Charlotte, Charlotte, North Carolina, USA; Florida Southern College, Biology Department, Lakeland, Florida, USA; Current address: Gregor Mendel Institute of Molecular Plant Biology, Austrian Academy of Sciences, Vienna, Austria.; Faculty of Life Sciences, Bar-Ilan University, Ramat Gan, Israel; The Multidisciplinary Brain Research Center, Bar-Ilan University, Ramat Gan, Israel

## Abstract

The ability of an animal to effectively capture prey and defend against predators is pivotal for its survival. Venom, a mixture of many toxin proteins, shapes predator-prey interactions. Here, we use the sea anemone *Nematostella vectensis* to test how toxin genotypes impact predator-prey interactions. We developed a new genetic manipulation tool which significantly reduces both RNA and protein levels of Nv1, a major neurotoxin. In concert we recently discovered a native population of *Nematostella* that has lost Nv1.We demonstrate that these anemones lacking Nv1, have reduced ability to defend themselves against grass shrimp, a native predator. Additionally, secreted Nv1 can act indirectly in defense by attracting mummichog fish, which are known to prey on grass shrimp. This unravels a tritrophic interaction acting in animal defense at the molecular level. Additionally, our work reveals an evolutionary tradeoff, as the reduction of Nv1 levels causes faster growth and increased sexual and asexual reproductive rates.

## Background

Capturing prey and defending against predators are vital tasks required for animal survival (*1*). During evolution, animals effectively optimize their predator and prey interactions while also balancing other traits essential for maintaining fitness such as gamete production and growth. As resources are limited, there is often a tradeoff between the energy an organism invests into each trait, and this tradeoff leads to a diversity of phenotypes we observe at the individual, population and species level (*2*). Uncovering the evolutionary tradeoff between metabolically costly traits is therefore essential to understand its impact on the phenotype and fitness of the organism. While understanding the genetic basis of a trait allows us to unravel the molecular processes selection acts driving phenotypic changes, connecting the genotype-phenotype link is challenging as most traits are shaped by many genes which often have multiple functions.

Venom is a trait involved in predator-prey interactions and is known to evolve under strong selection pressures (*3*). Venomous animals deploy a complex mixture of molecules to cause a physiological imbalance in another animal (*3*). A major component of venom is toxic proteins, known as toxins, which are directly encoded by the venomous animal’s genome. These toxin-encoding genes have been shown to evolve under strong selective pressure at both the sequence level and the gene expression level (*4–6*), likely due to the high metabolic cost of producing venom (*7, 8*). Furthermore, toxins are directly involved in predator-prey interactions and have been shown to have specialized functional roles. Striking examples of this specialized role of venom has been shown in multiple animals including scorpions, cone snails and assassin bugs, which can also dynamically shift their venom profile in response to predation or defense-evoked stimulation (*9–11*). Because of these factors, venom is an excellent system to understand functional adaptations at the molecular level by connecting the toxin genotype to the ecological phenotype, such as for predation or defense.

The sea anemone *Nematostella vectensis (***Fig 1A)** is arguably the most developed venomous model system, being amendable to genetic manipulation (*12*) and having much of the venom previously characterized (*13*). In *Nematostella*, Nv1 is a neurotoxin that modulates sodium channels and is highly lethal to various arthropods (*14, 15*). Nv1 is also the dominant toxin in adult venom, being produced in massive amounts and is among the most abundant proteins in the entire proteome (*5, 14, 16*). The abundance of protein is due to the Nv1 loci having more than 10 copies in a tandem array that all produce the same mature protein (*5, 17*). Our previous work revealed that Nv1 has undergone significant variation in copy number across populations (*5*). This is most striking in the population originating from Florida (FL), which has undergone a contraction event resulting in only a single diploid copy **(Fig 1B).** This contraction event resulted in a significant reduction of Nv1 at the transcription level compared to other populations, including the North Carolina (NC) population which have close genetic distance (*5*), as well as the Maryland (MD) population, which is the source of the lab strain **(Fig 1C, Table S1; ANOVA *P*-value = 0.0001)**. At the protein level, Nv1 is undetectable in FL compared to being among the top five most abundant protein from NC *Nematostella* (*5*). We find that the loss of Nv1 in FL is not an anecdotal phenomenon, with additional sampling of *Nematostella* 12 km from the original site, revealing that all individuals (n=10) have 0-2 Nv1 diploid copies (**Table S1**). While this work characterized the molecular signature that underlies Nv1 variation across populations, how this variation impacts key life history characteristics such as predator-prey interactions and reproduction, remains to be tested. Such insights would allow us to connect how changes in a genotype can impact the fitness and potential tradeoffs for an animal.

**Fig. 1.**
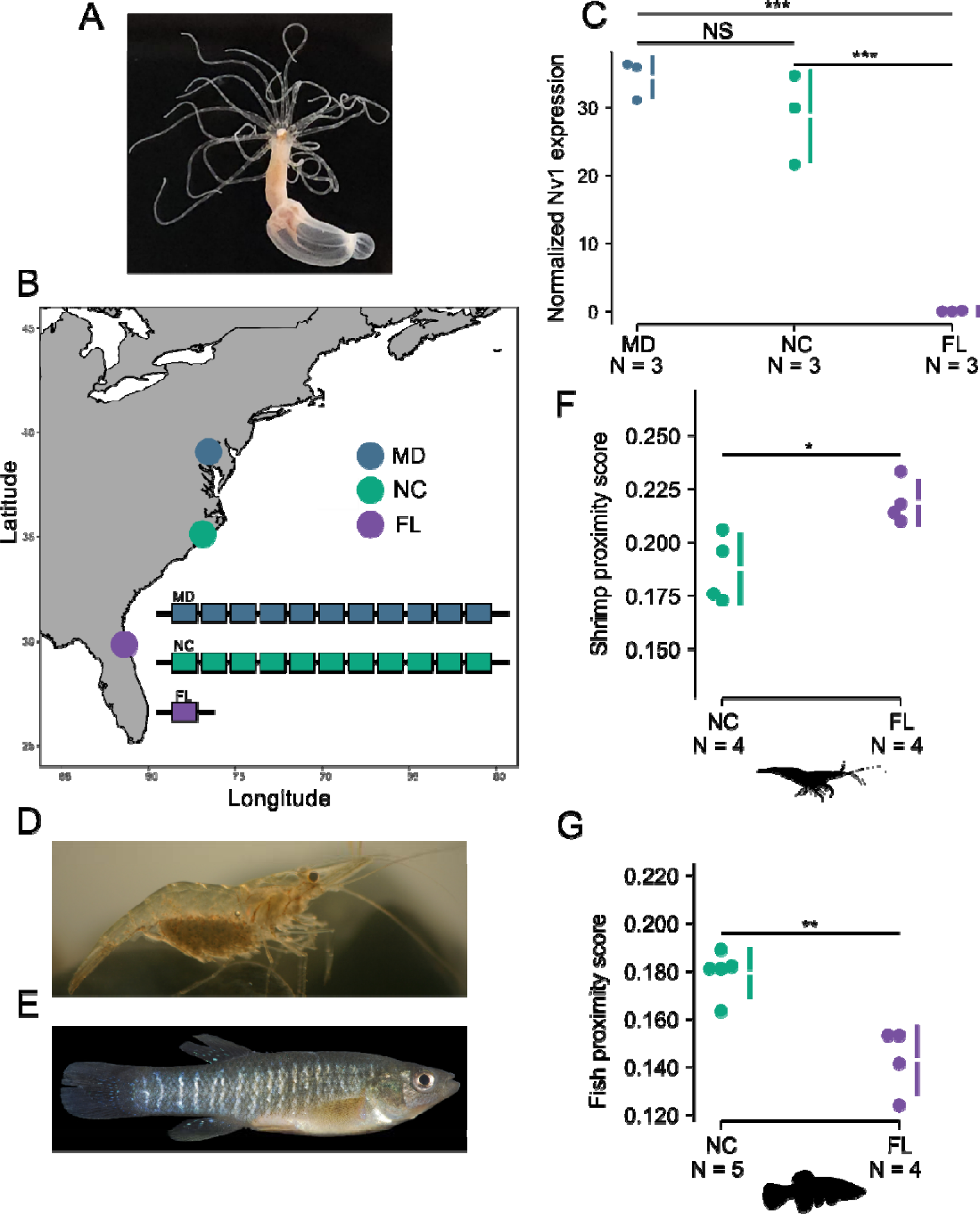
Population variation of Nv1 and its impact on defense against predators. A) *Nematostella vectensis*. B) Map showing the location of the different populations of *Nematostella* across North America. Florida (FL), Maryland (MD), North Carolina (NC). Inset, maximum haploid copy number of Nv1 reported in different *Nematostella* populations. C) nCounter relative RNA expression levels of Nv1 from different populations. D) grass shrimp (*Palaemonetes pugio*). Picture taken from Wikimedia by Brian Gratwicke (2006). E) mummichog (*Fundulus heteroclitus*) picture taken from Wikimedia from the Smithsonian Environmental Research Center. F) Average weighted score of grass shrimp proximity to *Nematostella* from NC and FL. G) Average weighted score of mummichog proximity to *Nematostella* from NC and FL. 95% confidence intervals are indicated by the ends of the vertical error bars. * *P*-value < 0.05, ** *P*-value <0.01, *** *P*-value <0.001, NS not significant.

To unravel how a toxin genotype can impact venom phenotype and affect an individual’s fitness, we investigated the impact of losing a toxin at the organismal level using a combination of behavioral and organismal assays. We developed a novel genetic manipulation technique using transgenesis to constitutively knockdown all Nv1 copies and challenged *Nematostella* of this transgenic strain as well as those of the native population from FL to capture prey and defend themselves against predators. We find that the FL population and the transgenic knockdown line, which both have depleted Nv1 levels, phenocopy and have a significantly reduced capacity to defend themselves against native predatory shrimp compared to *Nematostella* with wild-type levels of Nv1. In contrast, when we test these animals against native fish known to predate *Nematostella* larvae, we find that *Nematostella* that produce wild-type amounts of Nv1 attract the fish. Further experiments reveal that indeed Nv1 is being continuously secreted by *Nematostella* into the water and is detected by the fish, suggesting that it is acting as an attractant. Finally, when testing the physiological impact of synthesizing the massive amounts of Nv1, we find that those with depleted Nv1 levels grow faster and are more reproductive, highlighting that Nv1 synthesis is implicated in a significant evolutionary tradeoff with key fitness-related characters.

## Results and discussion

### Population variation of Nv1 and its impact on interspecific interactions

We set out to experimentally test the impact of losing Nv1 in predator-prey interactions. We first tested the ability of *Nematostella* to defend itself against ecologically relevant predators (*16, 18, 19*), specifically grass shrimp (**Fig 1D**; *Palaemonetes pugio*) and mummichogs (**Fig 1E**; *Fundulus heteroclitus*). We hypothesized that FL *Nematostella* that have reduced levels of Nv1 would have greater difficulty in defending themselves against these known predators. To test this, we recorded the interactions of *Nematostella* and each predator independently in a small vessel and scored the location of the predator based on its distance. *Nematostella* individuals that received a low score were considered those that were able to defend themselves, whereas a high score reflected a reduced ability to defend themselves. Our results reveal that *Nematostella* originating from Florida have a reduced capacity to defend themselves against grass shrimps, receiving a higher score compared to *Nematostella* from NC (**Fig1 F, Table S2**; ***P*-value = 0.016**).

The current results obtained for the differences among *Nematostella* native populations correspond to results from venomous snakes which have also undergone significant expansion and contraction events of toxin gene families. A prominent example was shown in the venom of *Trimeresurus flavoviridis* found on Okinawa to have a divergent venom composition when compared with other surrounding islands (*20*). Snakes from Okinawa were found to have lost the highly active myotoxic phospholipase A2 (PLA2) and was correlated with differences in their prey items, with snakes from Okinawa feeding on frogs compared to snakes on surrounding islands that have myotoxic PLA2 and feed on rats (*20*). Further studies in other viperid snakes have also shown similar correlations between diet and venom in different populations of the same species or closely-related species (*21–23*). Furthermore, it has also been suggested in snakes that predation and not defense drives the evolution of the venom phenotype, however notable examples contradict this (*24*). For example, spitting venom from three different lineages of cobras have convergently evolved for defensive purposes and have similar venom profiles (*25*). This is strongly supporting that defense from predation is also an important selective force driving venom evolution in some cases. Nevertheless, a functional proof is lacking in the examples due to the limitations in the genetic toolkit available in these organisms, which is essential as it cannot be excluded that other local adaptations could underlie those differences.

The ecological significance of venom in defense is evident in *Nematostella*, as venom components are present throughout its life history, yet *Nematostella* is only predatory after it settles and metamorphoses to a polyp (*16, 26*). Taken together with our findings that the loss of Nv1, the most abundant adult venom component, is correlated with reduced ability for *Nematostella* to defend itself against grass shrimp, suggests that defensive venom has an essential ecological role. While this is a strong hint that the differences between populations are the result of a loss of Nv1 in Florida, other local adaptations could have occurred that underly these differences. We therefore aimed to genetically manipulate the lab strain of *Nematostella*, coming from Maryland, to target Nv1 specifically and test if it is essential for *Nematostella* to defend itself against predators.

### Unravelling the evolutionary tradeoff of Nv1 using genetic approaches

Since Nv1 is encoded by numerous gene copies the CRISPR-Cas9 system could not be used to delete it from the genome. Thus, to target the unique loci of Nv1 we successfully employed a novel approach to simultaneously and constitutively knockdown all copies of Nv1, referred to as KD. This approach relied on the cnidarian microRNA (miRNA) pathway, a specialized RNA interference (RNAi) pathway, that has been shown previously to effectively silence genes in *Nematostella* (*27, 28*). We generated two lines using meganuclease to randomly integrate a construct that included a short hairpin that targets Nv1 (KD) and a control line that included the same features as the KD construct except for the absence of the short hairpin (**Fig 2A**). Both constructs successfully integrated into the *Nematostella* genome and are able to pass on the transgene across generations (**Fig 2B)**. We further confirmed that the miRNA undergoes processing by the small RNA machinery by using an end-point stem-loop PCR (**Fig S1**). This approach yielded KD *Nematostella* that have a significant reduction of ∼90% of Nv1 at both the RNA and (**Fig 2C**, **Table S4, *P*-value = 0.004**) and protein levels (**Fig 2D, Table S5, *P*-value = 0.0027**). Overall, these findings support that we have successfully generated the first genetically modified animal with a reduced venom arsenal.

**Fig. 2.**
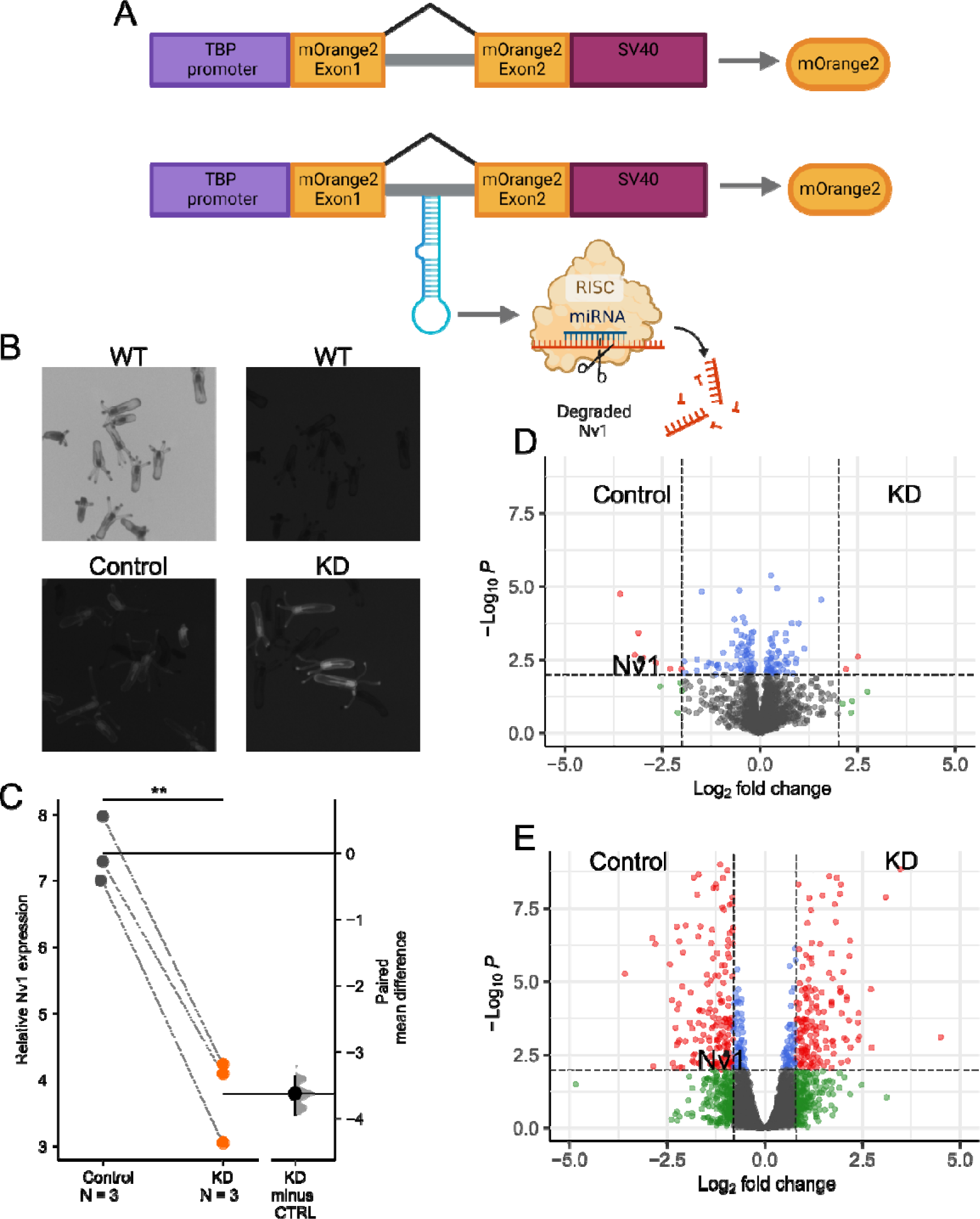
Characterization of the transgenic approach to constitutively knockdown Nv1. A) Schematic description of the vector used for transgenesis of both the control and knockdown line. A 2.9 kb TBP promoter was inserted upstream of the mOrange2 gene. Intron taken from NVE19315 (scaffold_507:85,148-86,273). The SV40 polyadenylation signal. KD construct also include a short hairpin designed to target Nv1. Created with BioRender.com. B) F2 animals showing successfully integration of both constructs compared non-fluorescent wild-type (WT) animals. C) RT-qPCR measuring the expression of Nv1. Plotted values are mean relative expression of Nv1 ΔCT comparing control to KD and represented as Gardner-Altman estimation plot. The mean difference is plotted on a floating axes on the right as a bootstrap sampling distribution. The mean difference is depicted as a dot; the 95% confidence interval is indicated by the ends of the vertical error bar. D) Volcano plot representing proteins of significantly different abundance, measured as label-free quantification (LFQ) intensity, between control and KD. Gray dots represent proteins that are not significant, blue dots are proteins with significant *P* value < 0.01, green dots are proteins with Log2 fold change of >2, red dots are proteins with significant *P*-value and Log2 fold change. E) Differentially expressed genes from 2-month-old animals. Differentially expressed genes (red) were defined by FDR < 0.01 and fold change ≥ 1.8. ** *P*-value <0.01, NS not significant.

### The role of Nv1 in defense

After the successful generation of *Nematostella* transgenic lines with reduced levels of Nv1, we aimed to test experimentally the role and significance of Nv1 specifically in interspecific interactions. We repeated our interspecific interaction assay with known predators, as described above, comparing our control with the KD line. Strikingly, when we test the interaction of the grass shrimp with transgenic *Nematostella* we see that the KD phenocopies the FL animals, with KD animals receiving a significantly higher score compared to control line animals (**Fig 3A, Table S8, Tukey HSD *P*-value = 0.007)**. This can be seen in video the footage, which shows the shrimp exhibiting a strong aversion to being touched by a control line animal (**Movie S1)** as well as exhibiting a clear avoidance swimming pattern around control line animals (**Movie S2)**. In contrast, shrimps exhibited a significantly reduced avoidance pattern often touching and walking over the KD animals (**Movie S3).** This supports that KD and FL animals have a reduced capacity to defend themselves against grass shrimp and provides strong evidence that Nv1 is essential in deterring this invertebrate predator. Furthermore, this confirms that a single venom component is essential for defense.

**Fig. 3.**
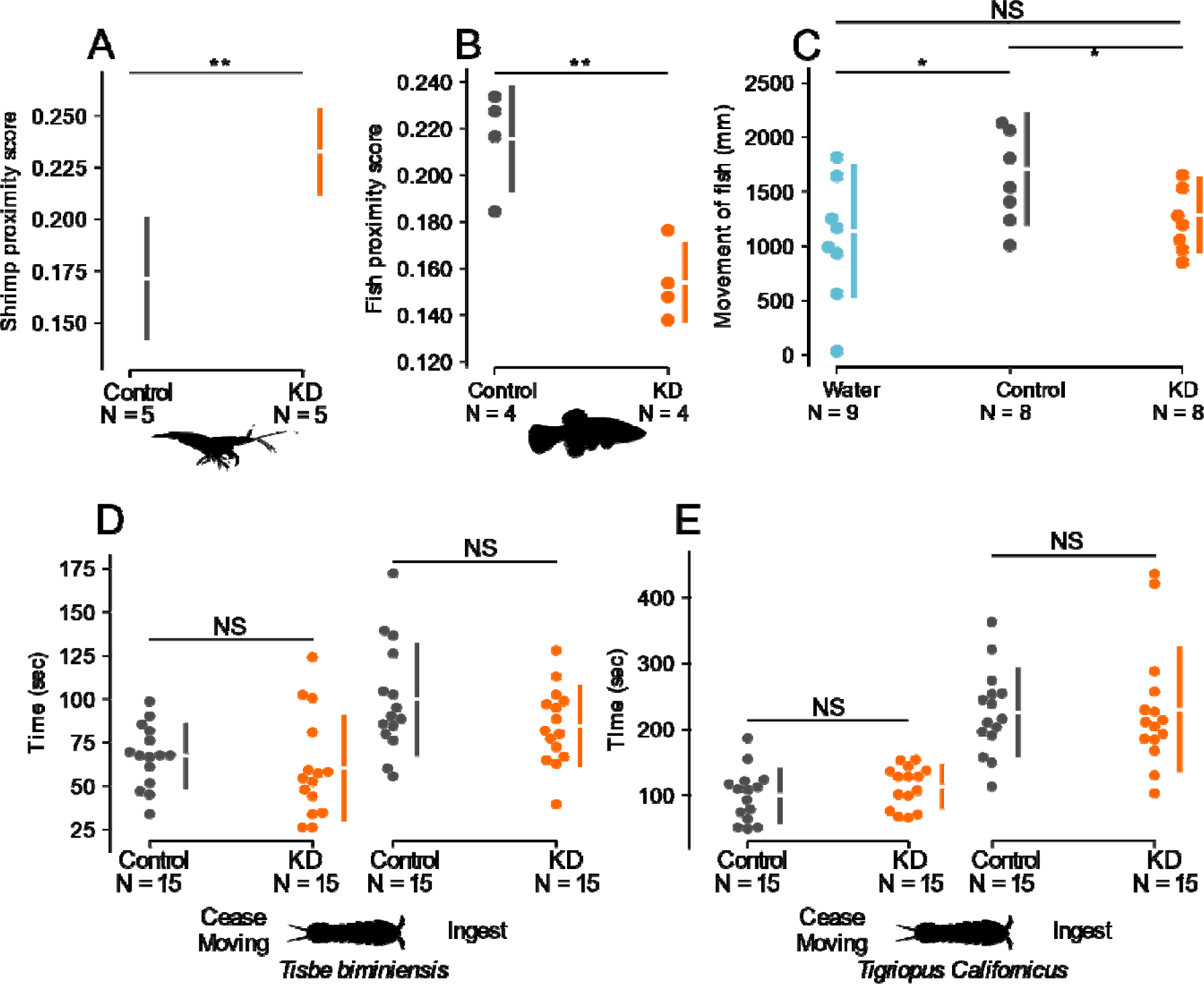
Impact of the depletion of Nv1 on defense and predation in *Nematostella*. A) Weighted score of grass shrimp proximity to *Nematostella* control and KD lines. 95% confidence intervals are indicated by the ends of the vertical error bars B) Weighted score of mummichog proximity to *Nematostella* control and KD lines. 95% confidence intervals are indicated by the ends of the vertical error bars. C) Tracking the movement of mummichogs in treated water coming from KD, control or 15 ‰ artificial sea water. D) Time taken for *Nematostella* to subdue movement and ingest *Tisbe biminiensis*. E) Time taken for *Nematostella* to subdue movement and ingest *Tigriopus californicus*. 95% confidence intervals are indicated by the ends of the vertical error bars. * *P*-value < 0.05, ** *P*-value <0.01, NS not significant.

In contrast, when we compare the interaction of the different *Nematostella* lines with mummichog fish, a known native predator of *Nematostella* larvae (*16, 19*), we report the inverse relationship **(Fig 1G and 3B),** compared to the grass shrimp. Specifically, we find that KD (**Fig 3B, Table S9, *P*-value = 0.004**) and FL (**Fig 1G, Table S3, *P*-value = 0.006**) *Nematostella* once again phenocopy each other as they both receive significantly lower scores, with mummichogs swimming in further proximity to these *Nematostella* compared to the control line and NC *Nematostella*, respectively. This led us to hypothesize that Nv1 is acting as a possible attractant as the mummichogs are swimming in greater proximity to *Nematostella* with high levels of Nv1 (control line and NC anemones). An alternative non-exclusive explanation is that the sharp reduction in Nv1 production in KD and FL animals leads to compensation by overexpression of other venom components that deter the mummichogs. However, both RNA-Seq experiments and proteomics failed to identify upregulation of other known toxins (**Fig 2 D and E, Table S5 and S6**).

We next suspected that for Nv1 to act as a possible attractant for the mummichogs, it would be required to be continually released into the water. This is supported by previous work which has shown that Nv1 is found in massive amounts in ectodermal gland cells (*16, 29*). Therefore, we first wanted to test if Nv1 protein can be captured in water incubated with *Nematostella*. To do this we incubated wild-type *Nematostella* overnight in 15 ‰ artificial sea water (ASW), and filtered it to remove any debris and nematocytes, referred to as treated water. The treated water was then concentrated using ultra-centrifugal filters and characterized with quantitative proteomics, revealing that indeed Nv1 is secreted into the water and is among the top 10 most abundant proteins (**Table S10**). The high levels of Nv1 released into the water would therefore allow other animals to encounter it without the need to directly contact the anemone.

We next tested if treated water alone, coming from control or KD lines, could impact mummichog behavior. To do this we incubated two *Nematostella* overnight in 6-well plates and again filtered to remove any debris and nematocytes. Mummichogs were then placed inside these wells with the treated water and their movement recorded (**Fig 3C, Table S11**). Strikingly, we find that mummichogs placed in water incubated with control line animals moved significantly more than the KD line (***P*-value = 0.03**) or filtered 15 ‰ ASW (***P*-value = 0.025**). This confirms that Nv1 is released into the water continuously and significantly affects mummichog behavior. Taken together with our previous results that mummichogs are attracted to animals producing wild-type levels of Nv1, it is likely that Nv1 acts to attract to mummichogs to *Nematostella*.

While this result was surprising, previous research by Kneib (1988) showed that when mummichogs, grass shrimp and *Nematostella* are all placed in the same tank over 10-weeks, *Nematostella* numbers increase. In contrast, Kneib (1988) found that *Nematostella* numbers decreased when only grass shrimp and *Nematostella* are placed together for 10-weeks (*30*). Additionally, there is evidence that mummichogs prey on grass shrimp (*30*). These results suggest that attracting mummichogs likely aids adult *Nematostella* in defending itself against grass shrimp. To understand this result in greater detail we performed additional behavioral assays in zebrafish as it is the arguably most well-established fish model system for behavioral studies (*31, 32*). Although mummichogs and zebrafish are distantly related, our aim was to test if some functional adaptation has occurred in mummichogs that allows it to detect low levels of Nv1 in the water or to dissect the behavior Nv1 has on fish at a greater granularity. To do this we tested the effect of treated water coming from control line and KD animals, at twice concentration used for the mummichogs, on 14-day old zebrafish (*Danio rerio*) and found no significant difference in their locomotive activity under alternating dark and light condition **(Fig S3, Table S12)**. This suggest that some coevolution may have occurred between *Nematostella* and the mummichogs that allows the latter to sense Nv1 at low concentrations and cues an attraction to the anemone.

The role of venom acting as a potential chemosensory cue in an aquatic environment has been previously reported. It was found that conotoxin-Tx VIA, a neurotoxin from the venom of a cone snail, triggers a specific warning response by a prey snail of the genus *Strombus,* causing them to initiate increased escape behavior locomotion (*33*). Taken together with our results, venom-derived peptides secreted into the aquatic environment may have a more significant impact on ecosystems beyond direct predator-prey interactions such as having additional indirect effects. To date, complex indirect interactions between plants, herbivores, and their natural predators, have been reported in detail, often referred to as tritrophic interactions, and found to be an integral part of all ecosystems (*34–36*). This work has mostly focused on the role of herbivore-induced plant volatiles, as the plants can attract predators and parasitoids to defend themselves against the herbivore (*34–36*). While these volatiles have been explored at the ecological and mechanistic perspective in plants, tritrophic interactions spanning across three levels of animals is scant and restricted to observational data, often referred to as indirect defense (*30*). Our findings are the first evidence that not only a venom component, but a single animal protein can dictate a tritrophic interaction required for defense. Overall, the role of venom in this complex interaction network is a novel discovery not previously reported. This is an exciting finding as the role of venom in chemical ecology has been largely overlooked and may have more systemic impacts than previously thought, especially in aquatic environments.

### The role of Nv1 on predation

We further tested the role of Nv1 in predation as intraspecific venom variation in snakes has been mostly linked to differences in diet (*21–23*). First, we explored the diet of native *Nematostella* across different populations using a metagenomic approach. We examined cytochrome c oxidase subunit 1 mitochondrial gene (CO1) sequences from the gut contents of *Nematostella* during the months of March, June and September, and in populations spanning the Atlantic coast of North America (Nova Scotia, Maine, New Hampshire, Massachusetts and North Carolina). The number of paired CO1 sequences varied tremendously among individuals at each location. For example, in March the Massachusetts population contained a single individual with 2,789 CO1 sequences (71% from that site alone), while another individual had no CO1 sequences recovered. All samples collected in March contained at least one individual lacking CO1 sequences, with the population from Nova Scotia having the most individuals lacking CO1 sequences (3/10, **Table S13**). This was even more pronounced in June and September, where the majority of individuals (43/60) across all locations returned no CO1 sequences and recovering a total of 139 and 10 CO1 sequences, respectively **(Table S13)**. Sequence abundance is indicative of the amount and time since recent predation events (*37*). Our results highlight that food abundances vary significantly across seasons and even among individuals, with periods of starvation being common. Additionally, we confirm that arthropods are a major food source for most populations (4/5, **Fig S4, Table S8**), *Nematostella* are largely opportunistic feeders with a large variety of potential prey items. This is consistent with previous work which focused on *Nematostella* from Nova Scotia also identified that arthropods are a major food source of *Nematostella*, but are ultimately opportunistic (*18*).

To determine the role and significance of Nv1 in capturing prey, we decided to focus on using copepods, which are among the top five most abundant arthropods found in the diet of *Nematostella* across all populations. Using two different copepods species, *Tigriopus californicus* and *Tisbe biminiensis*, we recorded the amount of time for the copepods to cease moving after contact with *Nematostella* and the amount of time for *Nematostella* to ingest after first contact. Overall, we find no significant difference between the control and KD lines in both metrics of predation for both species of copepod **(Fig 3D and E, Table S14)**. We further tested the predation success of these lines in capturing young mummichogs, but again find no differences (**Table S15**). Taken together, these results suggest that Nv1 likely plays a minor role, if any, in capturing prey.

This is in agreement with the spatial distribution of Nv1, which has higher expression in the physa compared to tentacles, pharynx and mesenteries (*16*). The physa is located at the aboral pole of *Nematostella* and is typically burrowed into sediment, when stressed *Nematostella* will typically retract the tentacles leaving only the body wall and physa exposed. In contrast, tentacles, pharynx and mesenteries all likely play important roles in catching and digesting prey where Nv1 levels are found in lower abundances. Contrastingly, these body region are abundant with nematocytes (stinging cells) which inject Nv1-less venom using a harpoon like structure (*38, 39*). Our results are in accordance with the recent finding that nematocytes in *Nematostella* are functionally specialized for predation, preferentially stinging when exposed to prey signals (*40*). This finding suggests that *Nematostella* separates its venom at both the anatomical and cellular level on the axis of predation and defense.

The separation of venom components and its specialization in predation or defense has been the focus of a significant body of recent research. To date, this research has shown that different venomous animals can produce distinct venom profiles following specific stimuli related to predation or defense. This was shown convincingly in scorpions, cone snails and the assassin bug, where they each secreted distinct venom profiles with different activities upon specific stimuli, highlighting that the molecular process for the separation of venom profiles has repeatedly evolved (*9–11, 13*). While these works showed that different venom profiles with distinct activities are induced with specific stimuli, the functionality was never tested at the organismal level. Here we show experimentally that toxin expression is separated for functionally distinct roles.

### The impact of Nv1 production on physiology and reproduction

Next, we hypothesized that the production of Nv1 has a high metabolic cost and therefore would impact the fitness of the organism, especially when resources are limited. Nv1 is produced in massive amounts at the protein level (*16*) and previous work has shown that it is metabolically expensive to replenish the entire venom system of *Nematostella* after depletion (*8*). Among the genes that contributed a major fraction transcriptionally during venom replenishment, was Nv1, further supporting that it likely has a high metabolic cost to synthesize (*8*). Finally, we see that upon starvation of four weeks, Nv1 levels in wild-type animals are downregulated ∼95 % (**Fig S5, Table S16, *P*-value = 0.006)**, which suggests that Nv1 levels are metered when resources are limited. We, therefore, decided to test if producing Nv1 impacts the growth and reproduction of *Nematostella*.

First, we began a feeding trial in which animals from both the control and KD lines were fed an excessive amount of food in the form of crushed brine shrimps. Once the animals reached four weeks old, we measured the length of the animals and observed no significant difference (**Fig 4A**, **Table S17, *P*-value = 0.75**). Following this, we began starving animals from both lines to see if relative growth rates are impacted when resources are limited. After two weeks of starvation, both lines continued to grow, with KD animals growing significantly more than control lines (**Fig 4B**, **Table S17, *P*-value = 0.046**). This significant difference continued after three weeks of starvation (**Fig 4C**, **Table S17, *P*-value = 0.034**). However, after four weeks of starvation, no differences were observed (**Table S17, *P*-value = 0.17**). After four weeks of starvation, however, we observed an additional nine animals in the containers of the KD line that could have only resulted from asexual reproduction by budding (*41*). A shift in reproductive strategy in the KD lines is supported by our enrichment analysis performed using our RNA-seq results, identifying the upregulation of genes related to cell-cycle and mitosis in KD *Nematostella* (**Fig 2E, Table S7**). This suggests that KD animals are undergoing increased rates of mitosis, allowing for faster growth and increased budding.

**Fig. 4.**
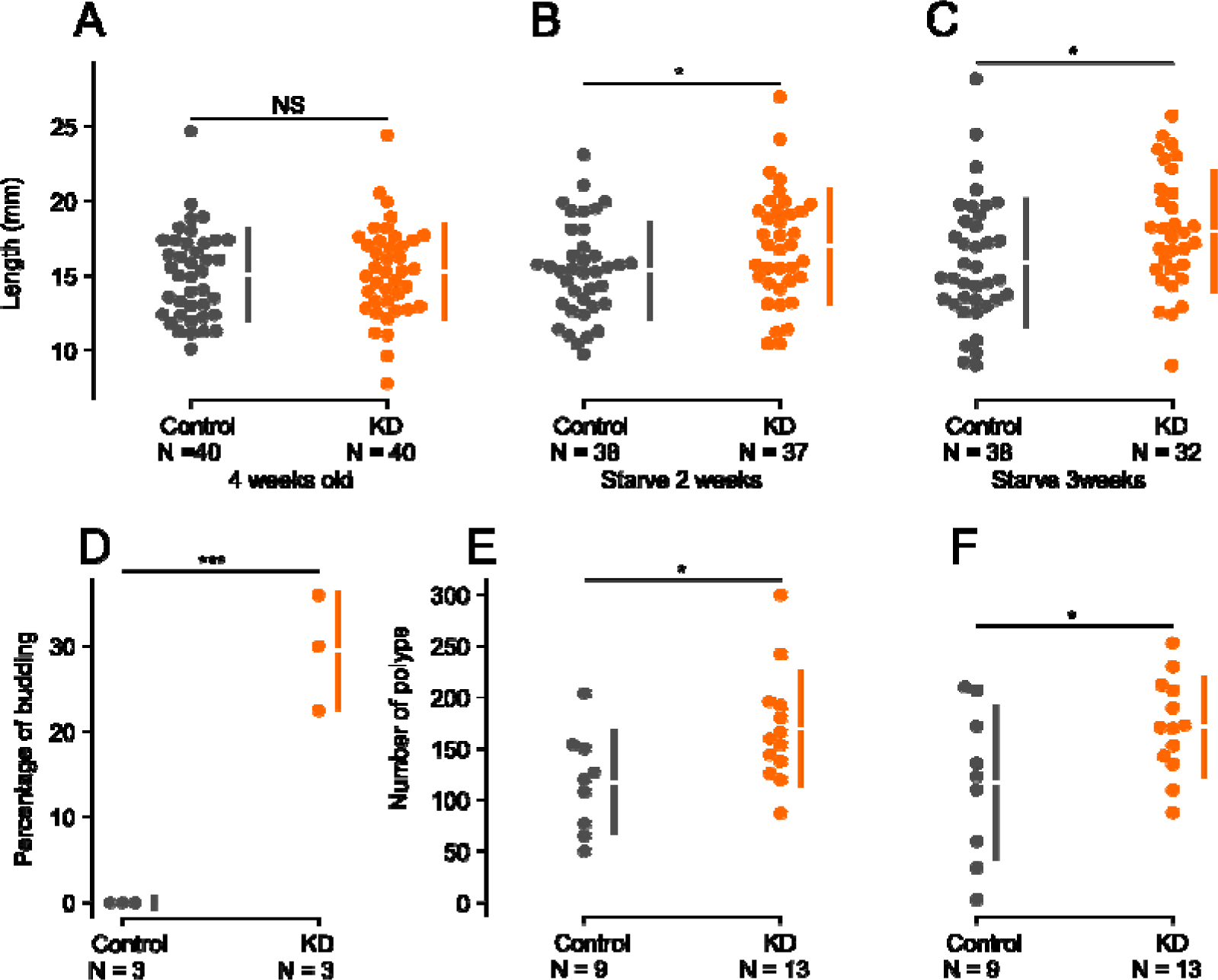
Impact of synthesizing of Nv1 on the physiology and reproduction of *Nematostella*. A) Length (mm) of 4-week-old *Nematostella* fed intensely. Length of 6-week-old *Nematostella* following 2 weeks of starvation. C) Length of 7-week-old *Nematostella* following 3 weeks of starvation. D) Percentage of *Nematostella* that went through asexual budding (n=3). E) Number of polyps developed from either control or KD transgenic eggs and fertilized with wild-type sperm. F) Number of successful polyps developed from either control or KD transgenic eggs following 2 weeks of starvation and fertilized with wild-type sperm. 95% confidence intervals are indicated by the ends of the vertical error bars. * P-value < 0.05, *** P-value <0.001, NS not significant.

In *Nematostella*, rapid growth is often associated with asexual reproduction via budding so we tested if KD animals asexually reproduce more. We find that the KD line asexually buds at significantly higher rates than the control line (**Fig 4D, Table S18, *P*-value= 0.0016**). Finally, we tested to see the impact of synthesizing Nv1 has on producing progeny through sexual reproduction. We find that females from the KD line produce significantly more polyps than the control line females (**Fig 4E, Table S19, *P*-value = 0.03**). We repeated this assay with the addition of starving the animals for two weeks as most animals in the field experience prolonged periods without food (**Table S13C**) and found that KD line still produces significantly more polyps than the control line females (**Fig 4F, Table S19, *P*-value = 0.04).** Taken together with the growth assay our results indicate that the production of a single venom component impacts the fitness of an animal. This is demonstrated by the fact that when Nv1 levels are reduced, animals can grow faster and reproduce more, both sexually and asexually.

This highlights that a significant evolutionary tradeoff is occurring in which the *Nematostella* populations must balance the amount of Nv1 they produce with their growth and reproduction rate. Tradeoffs between defense and growth has been reported extensively in plants and is commonly known as growth–defense tradeoffs in which plants adjust growth and defense based on external conditions (*42*). While some of these defensive traits are constitutive, such as development of a thick physical barrier (for example, bark), other defenses are highly inducible and respond only to presence of pests and result in reduced growth. In animals, tradeoffs between weaponry and reproduction have also been shown, such as in the horned beetle which balances the investment of horn-size used to obtain mates and testes growth (*43*). Unravelling the genetic basis that underlies these tradeoffs has been challenging as the traits are often complex and involve large gene networks or vast biosynthetic pathways. Furthermore, many examples of these tradeoffs are the result of ecological factors, such as pest presence or nutrient abundance, further challenging the ability to unravel the evolutionary history of these tradeoffs.

Here we have isolated a single heritable locus that underlies the balance between growth/reproduction and defense. Because of this resolution we can reconstruct and hypothesize the evolutionary history of Nv1 in *Nematostella*. Specifically, our findings when combined with previous work showing that Nv1 copy number varies significantly both within and among *Nematostella* populations (*5*), lead us to hypothesize that balancing selection is a major force in shaping the evolution of this loci. In our previous work we found that the Nv1 haplotype maintains heterogenous haplotypes and significant diversity in the copy number of Nv1 across populations ranging from 8-24 copies across Nova Scotia, Maine, New Hampshire, Massachusetts and Nort Carolina. In addition, variation within each population was also found, with no population reaching fixation, for example even in Florida some individuals have lost Nv1 completely, whereas others maintain two diploid copies. The role of balancing selection in venom evolution has been proposed in recent studies focusing on snakes where the maintenance of genetic diversity better explains the evolution of multiple venom gene families than directional positive selection (*44, 45*). We hypothesize that balancing selection may indeed be the major force driving the variation of Nv1 we observe. Heterogeneity and diversity of Nv1 copy number variation in each population helps to offset the costs producing too much or too little Nv1 related to growth/reproduction and defense. However, if selection pressures act intensely enough in favor of faster growth and reproduction at the expense of defense for multiple generations, this could explain the dramatic loss of Nv1 copies in the Florida population. Altogether, our work links venom genotype and phenotype to growth, reproduction and interspecific interactions via evolutionary tradeoff.

## Acknowledgments

Analysis was performed at the Stein Family mass spectrometry center in the Silberman Institute of Life Sciences, Hebrew University of Jerusalem. The authors are grateful to Dr. Dror Hawlena (The Hebrew University of Jerusalem) for help interpreting our interspecific interactions. The authors would also like to thank Dr. Shelby Rinehart (The University of Alabama) for guidance designing the behavioral assays. Finally, the authors would like to thank Dr Michal Bronstein and Mrs Adi Turjeman (The Center for Genomic Technologies, The Hebrew University of Jerusalem) for their help with high-throughput sequencing and Dr. Hanan Schoffman (Mass Spectrometry Core Facility, The Hebrew University of Jerusalem) for his help with proteomics. Species silhouettes were sourced from PhyloPic. grass shrimp (*Palaemonetes pugio*) picture taken from Wikimedia by Brian Gratwicke (2006). Mummichog (*Fundulus heteroclitus*) picture taken from Wikimedia from the Smithsonian Environmental Research Center. *Nemaotstella vectensis* picture taken by Yael Admoni.

## Funding

Lady Davis fellowship to J.M.S

Israel Science Foundation grant 636/21 (Y.M)

incentive funding from the CIPHER Center at UNC Charlotte (A.M.R)

Binational Science Foundation program with the National Science Foundation grants 1536530 and 2020669 (Y.M., A.M.R.)

German-Israeli Foundation for Scientific Research and Development (GIF Nexus, grant No. I-529-416.3-2021 to L.A.)

## Author contributions

Conceptualization: JMS, AMR, YM

Methodology: JMD, SB, JM, AJB, AF, RA, RR, JS

Investigation: JMS, SB, JM, LA, AMR, YM

Visualization: JMS

Funding acquisition: AMR, YM

Supervision: JMS, LA, AMR, YM

Writing – original draft: JMS, YM

Writing – review & editing: All authors

## Competing interests

Authors declare that they have no competing interests.

## Data and materials availability

The mass spectrometry proteomics data have been deposited to the ProteomeXchange Consortium via the PRIDE (46) partner repository with the dataset identifier PXD043447. Raw sequencing data for *Nematostella* control and KD lines are available NCBI SRA database for transcriptomics (BioProject PRJNA984949) and the metagenomics raw sequencing data is available NCBI SRA (SRR25074850 - SRR25074860).

## Supplementary Materials

Materials and Methods

Supplementary Text

Figs. S1 to S6

Tables S1 to S19

References (*1–29*)

Movies S1 to S3

## Notes

### Competing Interest Statement

The authors have declared no competing interest.

