## Supplementary Figures and Methods for "Venom tradeoff shapes interspecific interactions, physiology and reproduction"

**Affiliations:**

**The PDF file includes:**

Materials and Methods

Supplementary Text

Figs. S1 to S6

Tables S1 to S19

References (*1–29*)

**Other Supplementary Materials for this manuscript include the following:**

Movies S1 to S3

### Method

#### *Nv1 copy number variation across Nematostella populations*

We used quantitative PCR to determine the number of Nv1 copies in individuals collected from a new location in Florida, using methods previously published (*1*). Briefly, DNA for anemones from Nova Scotia, the original Florida location, and 10 individuals from a new Florida location was isolated with a DNeasy Blood & Tissue kit (Qiagen) using the manufacturer's protocol. Primers were used for Nv1 and Catalase (*1*). Amplifications were performed in a Bio-Rad CFX96 Touch Real-Time PCR Detection System using the Luna Universal Probe Mix (NEB). We performed three technical replicates of Catalase and Nv1 for each sample. Our plate setup contained triplicate reactions of a reference sample of known copy number from the original Florida population (diploid copy number = 1; derived from genome assembly). The diploid copy number was estimated using the ΔΔCt approach with Catalase as the single-copy control gene and the original Floridian sample of known copy number as our reference sample.

#### *Transgenesis*

Two synthetic gene fragments (gBlock, IDT, Belgium) were designed to target the unique loci of Nv1. This included the mOrange2 sequence (*2*), and included sequence taken from the intron that houses miR-2030 (*3*) in *Nematostella* (scaffold_507:85,148-86,273). For our KD line, an artificial miRNA was also included pre-miRNA sequence of Nve-miR-2022 (*3*) and changed the mature miRNA sequence to fully match the Nv1 transcript, an approach previously described to generate efficient knockdowns (*4*). Following digestion with restriction enzymes, PCR fragments were ligated to a pER242 (*5*) vector containing a TBP promoter previously proved to drive ubiquitous expression in *Nematostella* (*6*) and SV40 polyadenylation signal. Plasmids were transformed into the *Escherichia coli* DH5α (New England Biolabs, USA) strain and outsourced for Sanger sequencing (HyLabs, Israel). Each plasmid was subsequently injected into *Nematostella* zygotes along with the yeast meganuclease I-SceI (New England Biolabs) enable genomic integration (*5*, *7*). Transgenic animals were visualized under an SMZ18 stereomicroscope equipped with a DS-Qi2 camera (Nikon, Japan) and positive animals were reared to the adult stage. At approximately 4 months old F0 individuals were induced for gametes and crossed with wild-type animals to generate F1 heterozygotes. Multiple positive F1 adults for both lines were then crossed with wild-type animals, and these F2 animals were used for subsequent experiments.

#### *Knockdown validation*

To test the knockdown efficiency of our KD line vs control lines we performed qPCR and semi-quantitative proteomics. Since previous work has shown that Nv1 expression levels are highest in adult physa we chose to test the knockdown efficiency on this tissue type and developmental stage. First, adult animals (>3 months old) were anesthetized with MgCl2 and their physa was dissected.

##### *RNA extraction*

Total RNA was extracted (three replicates made from three individuals each) using TRIzol Reagent (Thermo Fisher Scientific, USA) and following the manufacturer's protocol. Samples were treated with 2 µl of Turbo DNAse (Thermo Fisher Scientific) and underwent an additional round of extraction using TRIzol Reagent (Thermo Fisher Scientific). A total of 500ng of RNA was reverse transcribed into cDNA using iScript cDNA Synthesis Kit (Bio-Rad, USA).

*qPCR*

Real-time qPCR was performed using Fast SYBR Green Master Mix (Thermo Fisher Scientific) on the StepOnePlus Real-Time PCR System v2.2 (Thermo Fisher Scientific). Nv1 expression levels were normalized to the NVE5273 gene (ΔCt = Ctreference gene − Ctgene of interest) which has been previously shown to have stable expression level across development and the relative expression calculated 2ΔΔCt method (*8*, *9*). The significance level was calculated by applying paired two-tailed Student's t-test to ΔCt values for each of the pairwise comparison between control and KD lines. Plots were made using estimation stats (*10*)

For the quantification of the miRNA in the KD line we designed the stem-loop primer (*11*, *12*). Next, we used 100 ng of total RNA for cDNA preparation using the SuperScript III Reverse Transcriptase (Thermo Fisher Scientific) and used random primers as a negative control or stem-loop. The presence of the miRNA primers was determined by using end point PCR (*13*), using 2 µl of cDNA as template, miRNAs-specific forward primer and stem-loop-specific reverse primer and run the PCR at 94°C for 2 min, followed by 35 cycles of 94°C for 15 s and 60°C for 1 min (*12*).

*Sample preparation for MS analysis*

Dissected adult physa was also used for semi-quantitative MS/MS analysis and performed using adults (five replicates, each made of three individuals) from both control and KD lines. Samples were snap frozen and lysed using 8 M urea and 400 mM ammonium bicarbonate solution. Lysed samples were centrifuged (22,000 g, 20 min, 4°C) and the supernatant was collected. Protein concentrations were measured with BCA Protein Assay Kit (Thermo Fisher Scientific).

Protein samples were denatured and reduced in 100 µl of 8M urea, 10 mM DTT, and 25 mM Tris-HCl pH 8.0 for 30 min. Iodoacetamide (55 mM) was added and samples were incubated in the dark for 30 min. Urea was diluted by the addition of 8 volumes of 25 mM Tris-HCl pH 8.0 followed by the addition of sequencing-grade modified Trypsin (Promega Corp., USA) (0.4 µg/ sample) and incubated overnight at 37°C with gentle agitation. The resulting peptides were acidified by the addition of 30 µl 10% formic acid (FA) and desalted on C18 home-made stage tips. Peptides were eluted using 80% acetonitrile (ACN), 0.1% FA and dried. Peptides were reconstituted in 0.1% FA and Peptide concentration was determined byabsorbance at 280 nm.

*nanoLC-MS/MS analysis*

MS analysis was performed using a Q Exactive-HF mass spectrometer (Thermo Fisher Scientific) coupled on-line to a nanoflow UHPLC instrument, Ultimate 3000 Dionex (Thermo Fisher Scientific). From each sample, 0.35µg of peptides were injected. Peptides were separated over a 107 min acetonitrile gradient (4% to 50%) at a flow rate of 0.15 µl/min on a reverse phase 25-cm-long C18 column (75 µm ID, 2 µm, 100Å, Thermo PepMapRSLC). Survey scans (300–1,650 m/z, target value 3E6 charges, maximum ion injection time 20 ms) were acquired and followed by higher energy collisional dissociation (HCD) based fragmentation (normalized collision energy 27). A resolution of 60,000 was used for survey scans and up to 15 dynamically chosen most abundant precursor ions, with "peptide preferable" profile were fragmented (isolation window 1.6 m/z). The MS/MS scans were acquired at a resolution of 15,000 (target value 1E5 charges, maximum ion injection times 25 ms). Dynamic exclusion was 20 sec. Data was acquired using Xcalibur software (Thermo Fisher Scientific).

*MS data analysis*

Mass spectra data were processed using the MaxQuant computational platform, version 2.0.3.0. Peak lists were searched against an NVE FASTA sequence database with the addition of the mOrange2 protein sequence (https://figshare.com/articles/*Nematostella*_vectensis_transcriptome_and_gene_models_v2_0/807696). The search included cysteine carbamidomethylation as a fixed modification, N-terminal acetylation and oxidation of methionine as variable modifications and allowed up to two miscleavages. The 'match-between-runs' option was used for lysate samples. For the water sample, the "dependent peptides" option was used. Peptides with a length of at least seven amino acids were considered and the required FDR was set to 1% at the peptide and protein level. Relative protein quantification in MaxQuant was performed using the label-free quantification (LFQ) algorithm for lysate samples only (*14*). MaxLFQ allows accurate proteome-wide label-free quantification by delayed normalization and maximal peptide ratio extraction (*14*). Statistical analysis ($n = 5$) was performed using the Perseus statistical package, Version 1.6.2.2121. Only those proteins for which at least three valid LFQ values were obtained in at least one sample group were accepted and log2 transformed. Statistical analysis by Student's t test and permutation-based FDR ($P$-value <0.05). After the application of this filter, a random value was substituted for proteins for which LFQ could not be determined ("Imputation" function of Perseus). The imputed values were in the range of 10% of the median value of all the proteins in the sample and allowed the calculation of $P$-values.

*RNA-seq*

To further test the knockdown efficiency as well as if any other toxins are compensating for the reduction in Nv1 across the whole organism, we performed RNA sequencing and differential gene expression. RNA was extracted as described above with the exception of using juvenile whole organisms (5 individuals at 2 months old). Total RNA quality was assessed using Bioanalyzer Nanochip (Agilent, USA) with all samples having RNA Integrity Number (RIN) > 7. Libraries were generated from 100ng RNA using NEBNext Ultra II Directional RNA Library Prep Kit (New England Biolabs) for Illumina and sequenced using a 75 bp single-end sequencing on a NextSeq500 machine (Illumina, USA).

Raw reads were trimmed and quality filtered using Trimmomatic (*15*). Reads were mapped to a modified *Nematostella* transcriptome (https://figshare.com/articles/*Nematostella*_vectensis_transcriptome_and_gene_models_v2_0/8076) with Nv1 transcripts collapsed into a single contig. Mapping was performed using Bowtie2 (*16*), and the gene counts were quantified using RSEM (*17*). Differential expression analyses were performed using scripts from Trinity (*18*) using both DESeq v2.139 (*19*) and edgeR v3.167 (*20*). Differentially expressed genes were defined by FDR < 0.01 and log2 fold change ≥ 0.8. Genes identified by both methods were considered as differentially expressed. Biological replicates were quality checked for batch effect using sample correlation and principal component analysis. Volcano plots were generated using EnhancedVolcano (https://github.com/kevinblighe/EnhancedVolcano). Transcriptome was annotated using BLASTx against the swiss-prot database (accessed 01/04/23) and enriched GO groups identified using GOseq (*21*) Bioconductor package implemented in the in-built Trinity pipeline (*18*). An FDR cut-off of 0.05 was considered significant for the enriched or depleted GO terms.

#### *Interspecific interactions*

*Animal care*

*Nematostella* embryos, larvae, and juveniles were grown in the dark at 22°C in 15‰ artificial seawater (ASW), whereas polyps were grown at 18°C. All polyps were fed with *Artemia salina* nauplii three times a week. The induction of gamete spawning was performed as previously described (*22*). The gelatinous egg sack was removed using 3% L-cysteine (Merck Millipore) and followed by microinjection of the plasmids. All *Nematostella* individuals used in this study belonged to the common lab strain originating from Rhode River MD (*23*)

Fertilized mummichogs (*Fundulus heteroclitus)* eggs from Kings Creek, VA (37°18 16.2"N 76°24 58.9"W) and Scorton Creek, MA (41°43'52.1"N 70°24'51.3"W) were kindly provided by Dr. Rafael Trevisan (Duke University) and Diane Nacci (Environmental Protection Agency), respectively. They were kept in 15 ‰ ASW at room temperature until hatching (around 2–3 weeks) and used immediately for behavioral analyses. Experiments on mummichogs were performed under permit no. 17–018 granted by the Institutional Animal Care and Use Committee (IACUC) at the University of North Carolina at Charlotte (IACUC-22-041 (UNC Charlotte) according to ethical regulations of Office of Laboratory Animal Welfare (National Institutes of Health, USA).

The first batch of grass shrimps (*Palaemonetes pugio*) was collected at an estuary near Georgetown, SC (33°21'01.0"N 79°11'26.1"W). Animals were transported to the lab and kept in 15‰ ASW in recirculating aquaria until use. Interaction experiments were conducted in 15‰ ASW containers. Grass shrimps were fed every day with TetraMin tropical fish food (Tetra Holding, USA). Mummichogs were fed twice daily with *Artemia* reared in the lab.

*Nematostella interactions with predators.*

To test the interaction of grass shrimp with *Nematostella*, all animals were starved for 48 hours before the experiment. Grass shrimp and *Nematostella* from different lines were placed in a container (27.5cm in length, 8cm in width, and 9cm in height) with 1000-1200 mls of 15‰ ASW and onyx sand premium natural substrate (Seachem, USA) and separated by a divider for 30 minutes before recording the experiment for 15 minutes using Moticam 580 (Motic, China). Grass shrimp were gently introduced, and the recording started. Videos were exported and a 15-grid key was overlayed on top of the recording using Adobe Premiere Pro. Scoring was performed based on grid locations of grass shrimp every 10 seconds (**Fig S6**). For grass shrimp experiments we examined multiple transgenic lines. This included lab strain wild-type, TBP::mCherry, control, and KD lines. One-way ANOVA and Tukey HSD post hoc analyses were performed to determine significance in a pairwise manner. No differences were observed between WT, mCherry, and control lines. Experiments comparing control and KD lines did not include WT and mCherry lines. Grass shrimp interactions were also performed for the comparison of NC and FL animals and were analyzed using a two-tailed Student's t-test. Interspecific interactions between *Nematostella*-mummichog were also performed as above with the exception of a 15-minute acclimation, a 10-

minute recording time, and the use of a different substrate (White Coral Sand, Nature's Ocean Premium Marine Substrates, USA) to allow better visualization of the mummichogs.

*Exposure of young mummichogs to Nematostella treated water behavioral assay*

To test if Nv1 is secreted into the water and affects fish behavior, we performed an additional behavioral analysis of mummichogs' movement. First, to detect if Nv1 is being secreted into the water, we performed an MS/MS analysis on water incubated with *Nematostella* overnight. Twelve wild-type females were placed in 42 ml of 15‰ ASWovernight (16 hours). Water was then collected and filtered using a 0.22 µm filter (Merck Millipore, USA) to remove any debris and released nematocytes. Filtered treated water was then concentrated first using Amicon Ultra-15 Centrifugal Filter Unit 3kda (Merck Millipore) at 12°C and 2,686 g to a final volume of 0.5 ml. We further concentrated this down to 40 µl using Amicon Ultra-0.5 Centrifugal Filter Unit (Merck Millipore) at 12°C and 4,000 g and submitted it for MS analysis.

MS analysis was performed using a Q Exactive-Plus mass spectrometer (Thermo Fisher Scientific) coupled on-line to a nanoflow UHPLC instrument, Ultimate 3000 Dionex (Thermo Fisher Scientific). Approximately 0.45µg of peptides were injected. Peptides were separated over a 52 min acetonitrile gradient (4% to 50%) at a flow rate of 0.15 µl/min on a reverse phase 25-cm-long C18 column (75 µm ID, 2 µm, 100Å, Thermo PepMapRSLC). Survey scans (380–2,000 m/z, target value 3E6 charges, maximum ion injection time 50 ms) were acquired and followed by higher energy collisional dissociation (HCD) based fragmentation (normalized collision energy 25). A resolution of 70,000 was used for survey scans and up to 15 dynamically chosen most abundant precursor ions, with "peptide preferable" profile were fragmented (isolation window 1.8 m/z). The MS/MS scans were acquired at a resolution of 17,500 (target value 1e-5 charges, maximum ion injection times 121 ms). Dynamic exclusion was 60 sec. Data was acquired using Xcalibur software (Thermo Fisher Scientific). MS data analysis was performed as described above.

Next, we performed a behavior analysis of mummichogs in treated water coming from either control or KD animals. Prior to the behavioral assay of mummichogs , KD and control anemones were starved for 48 hours prior to beginning water trials. The behavioral assays took place in 6-well plates where each well contained 7 ml of 15‰ ASW. Two anemones were placed in each well according to the trial taking place (KD or control) for 24 hours. Anemones were removed and the water was syringe filtered with a 0.22 µm Millex GP Filter unit (Merck Millipore). After filtration, we began an acclimation step in a new 6-well plate filled with 15‰ ASW. We created a transfer basket by modifying 40 µm easy strainers by cutting off the handles and placing a single strainer in each well. Young mummichogs were collected, and a single fish larva was placed in a transfer basket in the acclimation well plate. Fish larvae were acclimated in the room for 10 minutes and were then transferred to the experimental well plate that contained filtered anemone water in a DanioVision recording chamber (Noldus Information Technology, Netherlands). Video recordings were captured for 15 minutes, where recordings began prior to the transfer. We performed 9 replicates of fish exposed to KD anemone water and control anemone water, along with 10 replicates of a control that contained only 15‰ ASW (no exposure to anemones). We then used EthoVision XT 9 and XT 12 software (Noldus Information Technology) to track fish movement which was used in the analysis. Movement was then recorded using DanioVison (Noldus Information Technology) every 0.1 seconds, where movement < 0.1 mm was considered noise and removed. Distances were then used to perform a one-tail Student's t-test as previous work has shown that zebrafish move more when incubated with the recombinant Nv1 compared to Bovine Serum Albumin BSA (*24*). FDR corrections were performed to correct for multiple comparisons.

*Exposure of young zebrafish to Nematostella treated water behavioral assay*

To detect if secreted Nv1 affects other fish behavior, we performed an additional movement assay using zebrafish. First, 28 females were placed in 100 ml of 15‰ ASW overnight (16 hours). Water was then collected and filtered using a 0.22 µm filter (Merck Millipore) to remove any debris and released nematocytes. Filtered treated water was then concentrated 4x using Amicon Ultra-15 Centrifugal Filter Unit 3kda (Merck Millipore) at 12°C and 2,686 g to a final volume of 0.5 ml. We then performed buffer exchange and used E3 medium and repeated this process three additional times.

Wild-type adult zebrafish were intercrossed and their progeny were kept under light-dark (LD) cycle. At 14 days post-fertilization (dpf), the zebrafish were placed in a 48-well plate, alternating between control and KD water. The plates were placed in the DanioVision tracking system (Noldus Information Technology) and allowed to acclimate for 15 minutes before recording their activity. The light intensity in the tracking system was set at 70 LUX (25% in the operating software) for all experiments. To observe their responses to LD transitions, the zebrafish experienced three cycles of 30 minutes of light followed by 30 minutes of darkness. Each experiment involved four independent assays, which were recorded and analyzed using the EthoVision XT 9 and XT 12 software (Noldus Information Technology), as previously described (*25*). The data analysis for total activity was conducted according to previously described threshold parameters (*25*). Zebrafish protocol was reviewed and approved by the Bar-Ilan University Bioethics Committee.

*Nematostella interactions with prey.*

To analyze the gut contents of *Nematostella* we used whole-body genomic DNA extractions using a protocol described in our previous study(*1*). Total genomic DNA was extracted using the AllPrep DNA/RNA Kit (Qiagen, USA) for 10 or more individuals across five locations spanning the Atlantic coast of North America (Cresent Beach, Nova Scotia; Saco, Maine; Wallis Sand, New Hampshire; Sippewissett, Massachusetts; Ft. Fisher, North Carolina) during the months of March, June, and September in 2016. To target gut contents we used the primers (*26*) (LCO1490:GGTCAACAAATCATAAAGATATTGG and HC02198:TAAACTTCAGGGTGACCAAAAAATCA) to amplify the Cytochrome C Oxidase submit 1 (CO1) along with the adapter overhang for Nextera Indexing. Total genomic DNA extractions and PCR amplifications were conducted similarly to previously reported (*1*). Briefly, PCRs with modified adapters were performed using HiFi HotStart Ready Mix (Kappa Biosciences, Germany) with the following conditions: 95ºC - 3 min; 8 x (95ºC - 30s, 40ºC - 30s, 72ºC - 1 min), 72ºC - 5 min. Successful amplification was checked using the same approach, with an increase in cycles (35 x) to visually inspect via gel electrophoresis. Samples without visible bands were still included for sequencing to potentially sequence low abundance PCR Products. Libraries were prepared for sequencing using the MiSeq Reagent Kit v3 (600 cycles) (MS-102-3003) and indexed using the Nextera XT Index Kit V2 and sequenced alongside 5% PhiX. Overlapping reads were joined using BBMerge (*27*) (Version 38.84) and duplicate sequences were counted within Geneious Prime 2023.1.1.

A cytochrome c oxidase subunit 1 mitochondrial gene (CO1) database was constructed using sequences retrieved from NCBI Genbank using the Entrez Direct (Edirect) utility (accessed

5/10/23), specifically nucleotide sequences that contained the term cox1. MiSeq sequences were then searched against the custom CO1 database using blastn, with the top blast hit retained as a candidate sequence for each sampling locality. A series of custom python scripts (https://github.com/JasonMacrander/Stella_Venom) were used to identify sequence abundance and taxonomic diversity across sites.

Prey immobilization and consumption were compared between control and KD anemones with observational feeding experiments. We used two copepod species (*Tigriopus californicus* and *Tisbe biminiensis*) purchased from AlgaeBarn (USA) and differ in size but are representative of the small arthropods that compose a significant portion of the anemone's natural diet (*28*). *Nematostella vectensis* juveniles for these experiments resulted from crosses of multiple wild-type females with a transgenic male from either control or KD lines. Positive polyps were then picked 10 days after fertilization. Newly settled four-tentacle juveniles were fed freshly hatched *Artemia* nauplii for 2-3 weeks to increase their size to eight- to twelve-tentacle juveniles. This size was determined in preliminary experiments to be necessary for the efficient capture of these two copepod species. Smaller juveniles were capable of capturing copepods but encounter rates were too low for reliable comparisons of capture and consumption. In preparation for the observations, anemones were fed freshly hatched *Artemia* 48 hours prior to the experiment and then placed in new 15‰ ASW. Fifteen anemones from each line (KD and control) were placed individually in a single well of a flat bottom 96 well plate with 200 µl of seawater. A single copepod was introduced into the well and observed using a stereomicroscope (Leica M80, Germany). The time to immobilization was scored as the time until the copepod ceased movement. The time to consumption was measured as the time until the copepod was fully ingested into the mouth. Times were recorded to the nearest second. The recorder was blind to the treatment group at the time of experimentation and observation. Statistical comparisons for time to immobilization and time to consumption were completed using Student's t-test and a *P*-value of 0.05 was used to assess statistical significance.

We further tested the role of Nv1 in predation of a vertebrate. We placed mummichog larvae with *Nematostella* adults in a small environment to measure capture efficiency for the KD, control, FL, and NC individuals. To do this we used a 15 ml Falcon tube, which contained 2 ml of sediment and 6 ml of 15 ‰ ASW. Sea anemones and fish were allowed to first acclimate for 15 mins. Fish were then introduced into the tube with the sea anemone in sediment, and capture and consumption was recorded over 15 minutes. For each line, we had 10 replicates.

***Physiological and reproduction assays***

We next aimed to test the impact of synthesizing Nv1 on the physiology of *Nematostella*. To do this we first investigated the growth rate of animals with our control lines with wild-type Nv1 levels against our KD line with depleted Nv1 levels. Specifically, 40 positive animals per line were picked and grown in small plates with 20 animals per plate and grown at 25°C. Each plate of polyps was fed with 100 µl of highly concentrated artemia that was ground using a micro pestle in a randomized manner four times per week and water changed after a minimum of three hours till animals were four weeks old. Animals were visualized under the SMZ18 stereomicroscope equipped with a DS-Qi2 camera (Nikon) and measured using Fiji (*29*). Animals were then starved with water changes once a week and measured as above. All measurements were performed blinded and used to perform Student's t-test.

Next, we tested the difference in asexual reproduction rates between control and KD lines. To do this we tracked the number of polyps for one month at three different ages, ranging from 1 month old (n=40), 2 months old (n=25) and 3 months old (n=25). To test differences in sexual reproduction we sorted a cohort of 36 animals from both control and KD lines. Females were identified for both control (9) and KD (13) lines following induction and gamete identification. Animals were induced again 2 weeks later to get them in cycle and on the third induction the egg packages from control and KD lines were fertilized with wild-type sperm. Once the animals reached 2 weeks old polyps were anesthetized using $MgCl_2$ and counted. Animal care and polyp quantification was performed blinded.

To understand the impact of starvation on the expression of Nv1, we performed a qPCR analysis. Adult wild-type females were first acclimated to being fed three times a week for one month and grown in 6-well plates. After acclimation, 9 females were starved, while 9 were fed under the same conditions and after four weeks, physa was removed from all animals and snap frozen (3 replicates with 3 individuals per replicate). RNA, cDNA synthesis and qPCR analysis were performed as previously described.

**Fig. S1.**

2.5% agarose gel picture of miRNA. Lane 1 ladder, lane 2 control cDNA using random primers, lane 3 control cDNA using Nv1-mimiR stem-loop, lane 4 KD cDNA using random primers, lane 5 control cDNA using Nv1-mimiR stem-loop.

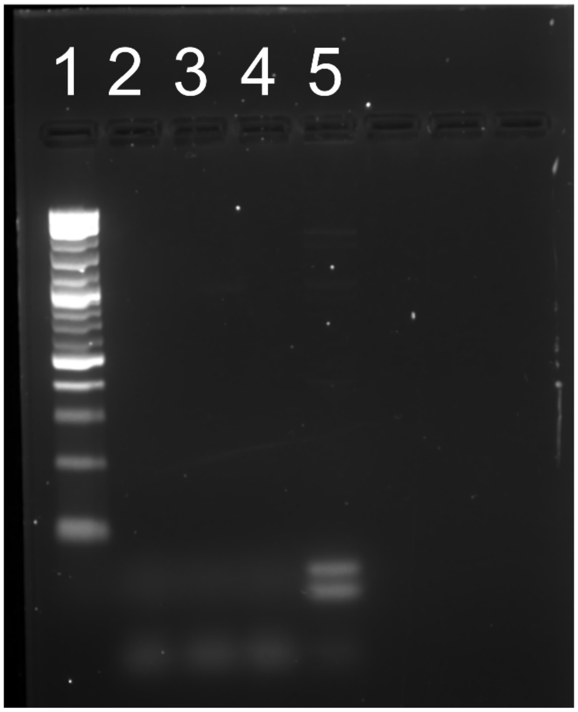

**Fig. S2.**

Average weighted score of grass shrimp proximity to *Nematostella* from different lines, including wild-type (WT), TBP::mCherry, control and KD lines. The letters above dot plots indicate the results of a Tukey–Kramer post hoc test, with lines showing significant differences ($P$-value < 0.05 for pairwise comparisons) unless they share the same letter. 95% confidence intervals are indicated by the ends of the vertical error bars.

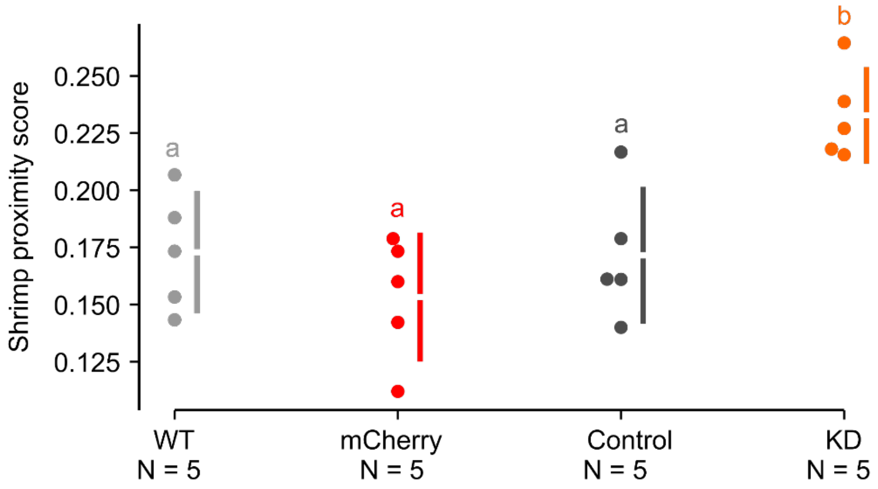

**Fig. S3.**

Tracking the movement of zebrafish. A) Average zebrafish movement (cm/min) over 180minutes in treated water coming from either KD or control *Nematostella* lines. A)95% confidence intervals are indicated by the ends of the vertical error bars. B) Average zebrafish movement for each minute, light and dark conditions with in dark conditions highlighted in grey.

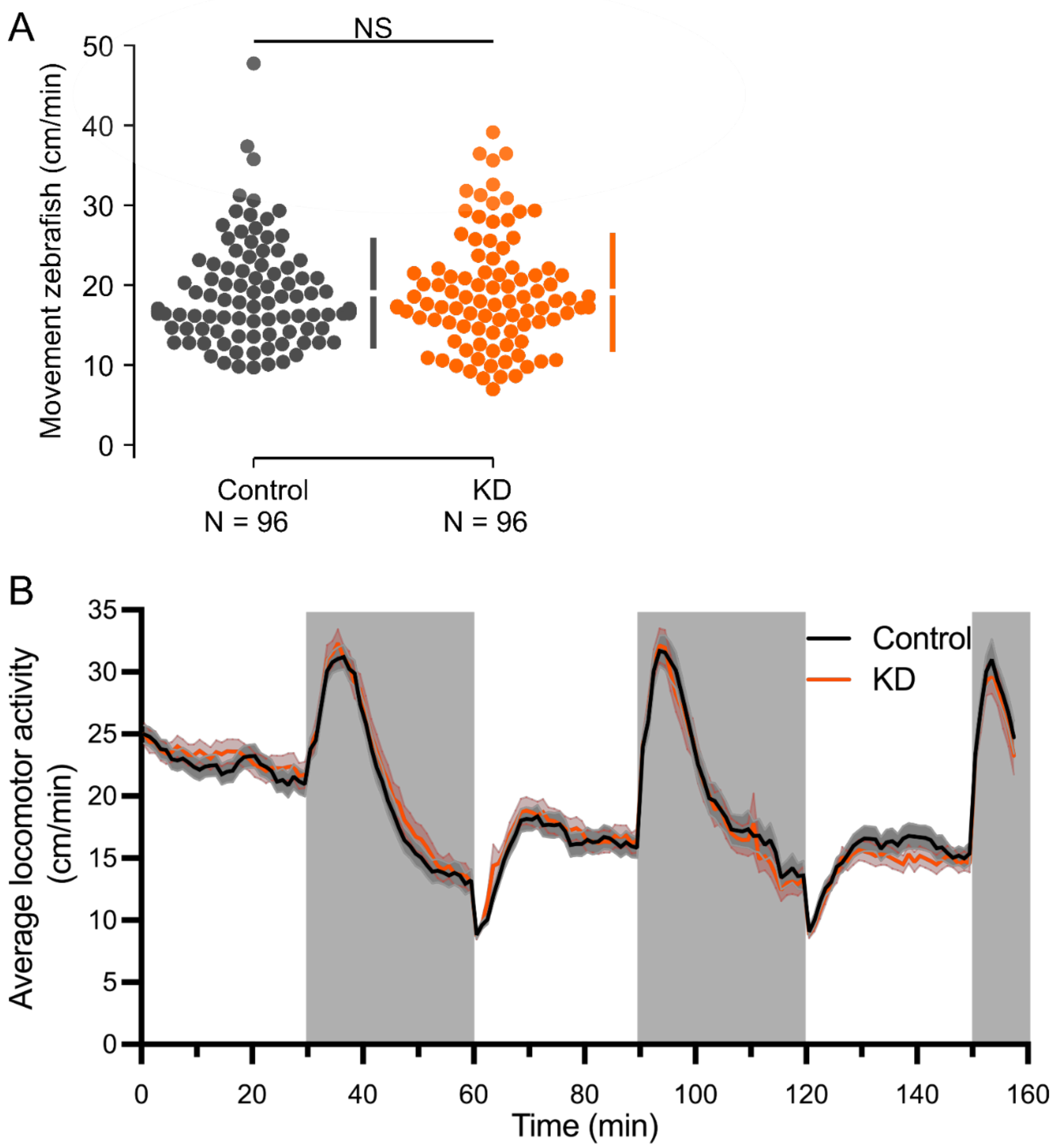

**Fig. S4.**

Metagenomics of gut content of *Nematostella* across different populations in March, June and September 2016. A) Percentage of reads from each population mapping at the phylum level, sequences mapping to Arthropod highlighted in black. B) Number of mapped reads in different populations across March, June and September. C) Cumulative number of reads log2 normalized from each population mapping to different arthropod species. North Carolina (NC), New Hampshire (NH), Massachusetts (MA), Nova Scotia (NS) and Maine (ME).

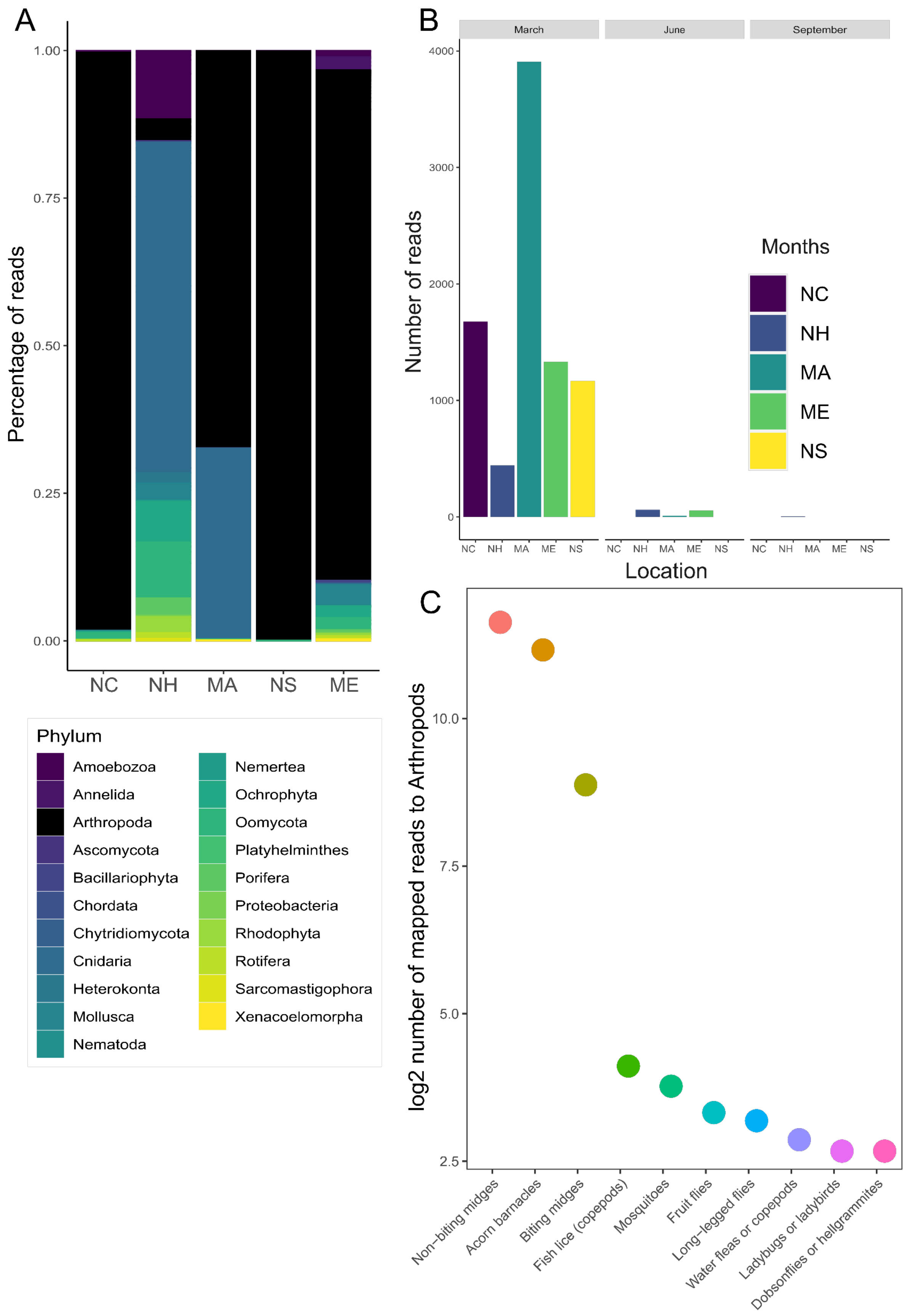
A
Percentage of reads
1.00
0.75
0.50
0.25
0.00
NC
NH
MA
NS
ME
Phylum
Amoebozoa
Annelida
Arthropoda
Ascomycota
Bacillariophyta
Chordata
Chytridiomycota
Cnidaria
Heterokonta
Mollusca
Nematoda
Nemertea
Ochrophyta
Oomycota
Platyhelminthes
Porifera
Proteobacteria
Rhodophyta
Rotifera
Sarcomastigophora
Xenacoelomorpha
B
March
June
September
Number of reads
4000
3000
2000
1000
0
NC NH MA ME NS
Location
Months
NC
NH
MA
ME
NS
C
log2 number of mapped reads to Arthropods
10.0
7.5
5.0
2.5
Non-biting midges
Acorn barnacles
Biting midges
Fish lice (copepods)
Mosquitoes
Fruit flies
Long-legged flies
Water fleas or copepods
Ladybugs or ladybirds
Dobsonflies or hellgrammites

**Fig. S5.**

RT-qPCR measuring the expression of Nv1. Plotted values are mean ΔCT comparing wild-type to starved *Nematostella*.. 95% confidence interval is indicated by the ends of the vertical error bar.

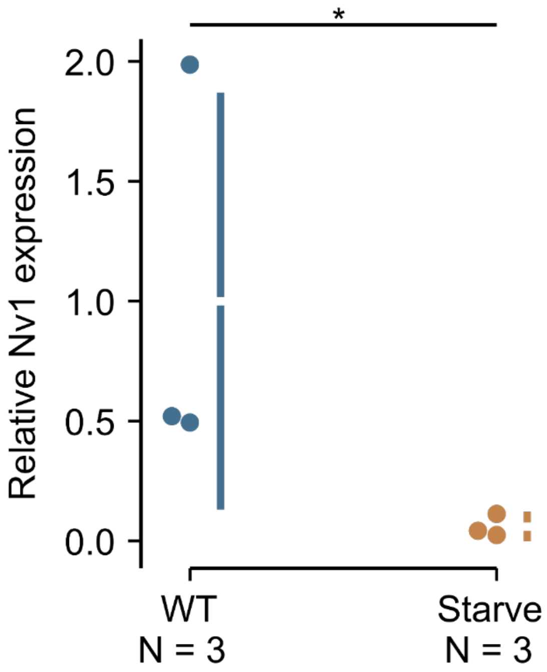

**Fig. S6.**

Setup for behavioral assay with the container used (A) and the scoring scheme used to quantify the proximity of predators to different lines of *Nematostella* (B).

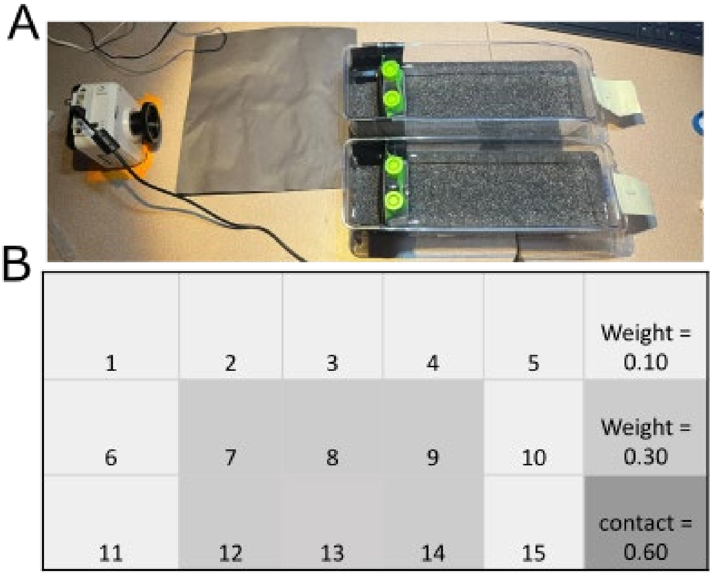

**Table S1.**

Meta-analysis nCounter data of Nv1 expression and housekeeping gene HKG4 across populations from different states of USA. Maryland (MD), North Carolina (NC), Florida (FL). A) Normalized nCounter data of Nv1 expression. C) ANOVA statistics included. B) Nv1 diploid copy number from Nova Scotia and two distinct locations from Florida.

**Table S2.**

Average weighted score of grass shrimp proximity to *Nematostella* from North Carolina (NC) and Florida (FL). Student's t-test (two-tail) also included.

**Table S3.**

Average weighted score of mummichog proximity to *Nematostella* against North Carolina (NC) and Florida (FL). Student's t-test (two-tail) also included.

**Table S4.**

A) Relative quantification (ΔCT) of Nv1 expression from control and KD lines. Student's t-test (two-tail) also included. B) Normalized relative quantification (ΔΔCT) of Nv1 expression from control and KD lines. C) Fold change difference Nv1 expression between control and KD lines. D) Knockdown percentage of Nv1 expression in KD line.

**Table S5.**

Perseus statistical analysis performed using semi-quantitative proteomic results from control and KD lines of *Nematostella vectensis*.

**Table S6.**

RNA-seq from control and KD lines of *Nematostella vectensis* with a log2foldchange of 0.8 and *P*-value 0.01. A) DeSeq2 result from genes upregulated in KD line. B) edgeR result from genes upregulated in KD line. C) DeSeq2 result from genes upregulated in control line. D) edgeR result from genes upregulated in control line

**Table S7.**

Go-term enrichment analysis using clusterProfile (A) and GoSeq (B)/

**Table S8.**

Average weighted score of grass shrimp proximity to *Nematostella* from wild-type (WT), TBP::mCherry (mCherry), control and KD lines. ANOVA statistics and Tukey HSD / Tukey Kramer included.

**Table S9.**

Average weighted score of mummichog proximity to *Nematostella* from control and KD line. Student's t-test (two-tail) also included.

**Table S10.**

Semi-quantitative proteomic result of treated water coming from wild-type *Nematostella*.

**Table S11.**

Average movement of mummichogs in water coming from untreated water (water), control and KD treated water. Student's t-test (one-tail) and FDR correction also included.

**Table S12.**

Average movement of zebrafish in water coming from control and KD treated water. Student's t-test (one-tail) also included.

**Table S13**

Metagenomics analysis of gut contents of *Nematostella* from different population. North Carolina (NC), New Hampshire (NH), Massachusetts (MA), Nova Scotia (NS) and Maine (ME). A) Number of CO1 sequences mapping back to animals in March 2016 at the phylum, order and family level. B) Total number of sequences from populations mapping in different months including March, June and September. C) Number of individuals from different populations that had no reads mapping to database.

**Table S14.**

A) Time (sec) taken for *Nematostella* to capture and ingest *Tisbe biminiensis in control and KD lines. B)* Time (sec) taken for *Nematostella* to capture and ingest *Tigriopus californicus in control and KD lines.*

**Table S15.**

Time (sec) taken for *Nematostella* to capture and ingest *mummichogs in NC, FL, control and KD lines.*

**Table S16.**

A) Relative quantification (ΔCT) of Nv1 expression in adult wild-type females (WT) and starved adult wild-type females. Student's t-test (two-tail) also included. B) Normalized relative quantification (ΔΔCT) of Nv1 expression from control and KD lines. C) Fold change difference Nv1 expression between control and KD lines. D) Knockdown percentage of Nv1 expression in KD line.

**Table S17.**

Growth assay of *Nematostella* following intensive feeding and measured (mm) at 4 weeks old followed by 2, 3 and 4 weeks of starvation for control and KD lines. Student's t-test (two-tail) also included.

**Table S18.**

Asexual reproduction rates from three independent observations measured as percentage of new polyps by original number of polyps for control and KD lines. Student's t-test (two-tail) also included.

**Table S19.**

A) Sexual reproduction rates measured as number of successful polyp development after 14dpf in control and KD lines. B) Sexual reproduction following 2 weeks of starvation in control and KD lines.

**Movie S1.**
Movie showing shrimp exhibiting severe recoil following a touch with a control animal.

**Movie S2.**
Movie showing shrimp exhibiting avoidance in swimming behavior when placed with control animal.

**Movie S3.**
Movie showing shrimp actively engaging with KD animal as well as some feeding behaviour.
